# Effects of chronic dietary hexavalent chromium on bioaccumulation and immune responses in the sea cucumber *Apostichopus japonicus*

**DOI:** 10.1101/2021.10.01.462835

**Authors:** Qixia Chan, Fuqiang Wang, Lidong Shi, Xue Ren, Tongjun Ren, Yuzhe Han

**Author notes:** Corresponding author (Fuqiang Wang).

## Abstract

Sea cucumbers *Apostichopus japonicus* (3.54 ± 0.01 g of wet weight) were exposed to five concentrations of dietary hexavalent chromium [0 (control), 100, 200, 400, and 800 mg Cr^6+^/kg dry weight] amended with K_2_Cr_2_O_7_ for 30 days. The bioaccumulation and immune responses [antioxidant enzymes: superoxide dismutase (SOD) and catalase (CAT); hydrolytic enzymes: acid phosphatase (ACP) and alkaline phosphatase (AKP)] of sea cucumbers were subsequently evaluated. This study found that the order of Cr accumulation in the experimental tissues was respiratory tree > intestine > body wall. Significantly lower SOD activities occurred in the 400 mg/kg group compared to that in the control group. Higher dietary Cr^6+^ exposure (400 and 800 mg Cr^6+^ /kg dry weight) did not negatively alter the CAT activities, but significantly inhibited CAT activities in 100 mg/kg group, compared to control group. ACP activities in groups 200, 400 and 800 mg/kg were significantly lower than those in control group, while no significant differences occurred in AKP activities among groups. The present study provides important information into the bioaccumulation and immune responses of the sea cucumber *A. japonicus* in response to chronic dietary Cr^6+^ exposure.

## 1. Introduction

Chronic contamination by heavy metals is one of the most important issues in aquatic environment because they invariably remain in the environment (Ashraf, 2005). In this context, chromium (Cr) has been well documented that causing health concerns and environmental pollution around the world (Dhal et al., 2013). Cr has different valence states (II–VI), and its chemical properties and toxicity vary with these valence states. Hexavalent chromium (Cr^6+^) is one of the most harmful pollutants listed by the United States′ Environmental Protection Agency (US EPA) (Jiang et al., 2019; Tang et al., 2021), because it induces negatively impacts at cellular, molecular and ecosystem (Olibet et al., 1995). Nowadays, Cr^6+^ is widely applied by many industries, such as dyeing, fungicide, ceramic glass and refining (Zahoor and Rehman, 2009). Serious pollution of Cr^6+^ occurred in aquatic environment due to improper disposal of these industrial wastes (Ashraf and Hanfiah., 2017; Tang et al., 2021). Using Cr^6+^ as the filling materials in the junkyard further makes it more easily to penetrate groundwater (Domingues et al., 2010). The concentrations of Cr^6+^ in aquatic environment (12 mg/L), for example, are much higher than the national standards in India (100 μg/L) (Sharma et al., 2012; Awasthi et al., 2018).

Sea cucumber *Apostichopus japonicus* is deposit animal (Ru et al., 2019) and like to habit seabed rich in algae and sea mud because of the abundant organic matters, such as benthic diatoms, bacteria and organic debris of animals and/or plants (Jiang et al., 2013; Yokoyama, 2013). The habits of sea cucumbers lead them to be more vulnerable to the heavy metals in sediments (Jiang et al., 2015). Sun et al. (2007) documented that Cr^6+^ in water displayed acute toxicity on *A. japonicus*, and the LC_50_ of 24 h, 48 h and 72 h were 31.974, 7.499 and 3.808 mg/L, respectively. Significantly lower growth rate occurred when sea cucumbers *A. japonicus* were fed Cr^6+^ diet after a short period (10 days) (Li et al., 2019). However, the bioaccumulation and immune of *A. japonicus*, which are the most important biological responses, have not been fully studied when they were exposed to chronic dietary Cr^6+^ exposure.

Here, sea cucumbers were chronic exposed to five concentrations of dietary Cr^6+^ [0 (control), 100, 200, 400, and 800 mg Cr^6+^ /kg dry weight] for 30 days to investigate: 1) How the dietary Cr^6+^ accumulates in different tissues (body wall, intestine, respiratory tree) of *A. japonicus*; 2) whether dietary Cr^6+^ affects the immune responses of *A. japonicus*.

## 2. Materials and methods

### 2.1 Experimental diets

Five formulated diets containing different levels of Cr^6+^ (0, 100, 200, 400, 800 mg Cr^6+^ /kg dry weight) amended with K_2_Cr_2_O_7_ (Wuxi Hongxing Chemical Industry Factory, Wuxi, China) were used in the present study (Table 1). The actual Cr concentrations were 19.87, 139.40, 243.70, 359.00, 930.60 mg Cr /kg dry weight, respectively. Dry ingredients were mingled (Xindou Food Machinery Factory, Jiangmen, China) with freshwater at a ratio of 600 g/kg (Wang et al., 2015). They then were shaped into pellets (1.88 mm of test diameter) using a pellet mill (Pinzheng Equipment Co. Ltd., Changzhou, China). All diets were subsequently dried at 40 °C in a hot air oven (Jinghong Test Equipment Co. Ltd., Shanghai, China) for 24 h and then stored at -20 °C for the following experiment.

**Table 1.** Ingredients composition and nutrient levels of experimental diets (g/kg)

| Ingredients | 0<br>mg/kg | 100<br>mg/kg | 200<br>mg/kg | 400<br>mg/kg | 800<br>mg/kg |
| --- | --- | --- | --- | --- | --- |
| Fish meal | 60 | 60 | 60 | 60 | 60 |
| Soybean meal | 100 | 100 | 100 | 100 | 100 |
| *Scagassum* | 300 | 300 | 300 | 300 | 300 |
| *Laminaria japonica* | 200 | 200 | 200 | 200 | 200 |
| Sea mud | 200 | 200 | 200 | 200 | 200 |
| Yeast | 40 | 40 | 40 | 40 | 40 |
| Flour | 50 | 50 | 50 | 50 | 50 |
| Vitamin mixture<sup>a</sup> | 10 | 10 | 10 | 10 | 10 |
| Mineral mixture<sup>b</sup> | 10 | 10 | 10 | 10 | 10 |
| Guar gum | 20 | 20 | 20 | 20 | 20 |
| Microcrystalline cellulose | 10 | 9.72 | 9.43 | 8.87 | 7.74 |
| K<sub>2</sub>Cr<sub>2</sub>O<sub>7</sub><sup>c</sup> | 0 | 0.28 | 0.57 | 1.13 | 2.26 |
| Total | 1000 | 1000 | 1000 | 1000 | 1000 |
| Analyzed nutrients (% in dry basis) |  |  |  |  |  |
| Moisture | 5.64 | 5.29 | 6.62 | 5.18 | 6.68 |
| Ash | 38.33 | 38.10 | 38.37 | 38.94 | 38.40 |
| Crude protein | 15.15 | 14.87 | 15.17 | 14.19 | 14.73 |
| Crude lipid | 6.07 | 5.58 | 5.42 | 6.17 | 5.26 |

<sup>a</sup> Vitamin mixture (per kilogram of premix): vitamin A, 1,000,000 IU; vitamin D<sub>3</sub>, 300,000 IU; vitamin E, 4,000 IU; vitamin K<sub>3</sub>, 1,000 mg; vitamin B<sub>1</sub>, 2,000 mg; vitamin B<sub>2</sub>, 1,500 mg; vitamin B<sub>6</sub>, 1,000 mg; vitamin B<sub>12</sub>, 5mg; nicotinic acid, 1,000 mg; vitamin C, 5,000 mg; Ca pantothenate, 5,000 mg; folic acid, 100 mg; inositol, 10,000 mg; carrierglucose and H<sub>2</sub>O ≤ 10%.
<sup>b</sup> Mineral mixture (0.025 mg/g of premix): NaCl, 107.79; MgSO<sub>4</sub>·7H<sub>2</sub>O, 380.02; NaHPO<sub>4</sub>·2H<sub>2</sub>O, 241.91; KH<sub>2</sub>PO<sub>4</sub>, 665.20; Ca(H<sub>2</sub>PO<sub>4</sub>)·2H<sub>2</sub>O, 376.70; Fe citrate, 82.38; Ca lactate, 907.10; Al(OH)<sub>3</sub>, 0.52; ZnSO<sub>4</sub>·7H<sub>2</sub>O, 9.90; CuSO<sub>4</sub>, 0.28; MnSO<sub>4</sub>·7H<sub>2</sub>O, 2.22; Ca(IO<sub>3</sub>)<sub>2</sub>, 0.42; CoSO<sub>4</sub>·H<sub>2</sub>O, 2.77.
<sup>c</sup> potassium dichromate (K<sub>2</sub>Cr<sub>2</sub>O<sub>7</sub>)

### 2.2 Sea cucumber and experimental design

Eight hundred juvenile sea cucumbers (∼3 g) were transported to Dalian Ocean University (121°56′ N, 38° 87′ E) from Dalian Yinhaima Aquatic Products Co., Ltd (121°54′ N, 39°38′ E) on 26 October 2020. They were maintained in a 250-L tank (length × width × height: 76 × 55 × 60 cm) with aeration and fed control diet for two weeks to acclimatized the lab environment. Sea cucumbers were fasted for 24 h before the experiment began on 6 November 2020.

Formulated diets with different concentrations of Cr^6+^ were the experimental factors. Each group has three replicates and the experiment duration lasted for 30 days (from 6 November 2020 to 5 December 2020). Twenty-five sea cucumbers (3.54 ± 0.01 g) were randomly selected and placed into each of 15 plastic tanks (length × width × height: 50 × 35 × 29 cm) with aeration according to the experimental design. Sea cucumbers were fed the corresponding diets at 3% of body weight once a day at 17:00 during the experiment (Wang et al., 2015; Li et al., 2018). These diets were not dissolved within 24 hours. One-half of seawater was exchanged every day. Water temperature, salinity, and pH were measured weekly by a portable water quality meter (HACH, HQ40D, USA). They were 14.73 ± 0.26 °C, 31.2 ± 0.13‰ and 7.92 ± 0.01, respectively. Five sea cucumbers from each tank were randomly chosen before they were starved for 24 h. The coelomic fluid, body wall, intestine, and respiratory tree of sea cucumbers were obtained and kept at -80 °C for the further analyses of enzyme activity of coelomic fluid and Cr concentrations in tissues (body wall, intestine and respiratory tree) (N = 3).

### 2.3 Growth performance

The final body weight (FBW), body weight gain (BWG), specific growth rate (SGR), and survival rate (SR) were assessed at the end of the experiment on 5 December 2020 (N = 3). Growth parameters were calculated according to Wang et al (2016): BWG (%) = 100 × (final mean body weight − initial mean body weight) / initial mean body weight; SGR (%/day) = (ln final mean body weight − ln initial mean body weight) / feeding days; SR (%) = 100 × (final sea cucumber number) / (initial sea cucumber number).

### 2.4 Chromium concentration in sea cucumber tissues

The Cr^6+^ concentration in experimental tissues was analyzed on days 10, 20 and 30 according to Kim and Kang (2016) as follows. The 65% HNO_3_ (v/v) was added to digest these tissues at 120 °C and the digested solution was further diluted with 2% HNO_3_ (v/v) to make ∼25 mL liquid following a filtration using a membrane filter (0.2 µm). The Cr concentration in these tissues was determined using atomic absorption spectrometer (Perkin Elmer Instruments Shanghai Co. Ltd. Shanghai, China) and expressed as mg/kg wet weight.

### 2.5 Biochemical analysis

The enzymes activities were measured at the end of the experiment following the kit introductions (Nanjing Jiancheng Bio-engineering Institute, Nanjing, China). The superoxide dismutase (SOD) was measured according to Marklund and Marklund (1974). The activities of catalase (CAT) were analyzed flowing Góth (1991). The levels of malondialdehyde (MDA) were quantified according to Scown et al. (2010). The activities of acid phosphatase (ACP) and alkaline phosphatase (AKP) were determined by colorimetric method (Malamy and Horecker, 1966).

### 2.6 Statistical analyses

The differences of SOD activities were analyzed using Kruskal-Wallis H. One-way ANOVA was performed to analyze the differences of all remaining data followed by the LSD multiple comparison. All data were expressed as mean ± S.E and analyzed using SPSS software version 19.0 (SPSS, Chicago, IL, USA). A value of *P* < 0.05 was considered significant.

## 3. Results

### 3.1 Growth performance

Significantly lower BWG and SGR occurred in 800 mg/kg group than those in control group after 30-day Cr^6+^ exposure (*P* < 0.05, Table 2). However, there was no significant difference in SR among groups (*P* > 0.05, Table 2).

**Table 2.** Effects of dietary Cr^6+^ on the growth performance of sea cucumbers *Apostichopus japonicus*.

| Index | 0 mg/kg | 100 mg/kg | 200 mg/kg | 400 mg/kg | 800 mg/kg |
| --- | --- | --- | --- | --- | --- |
| IBW (g) | 3.53±0.01 | 3.55±0.02 | 3.55±0.03 | 3.54±0.01 | 3.53±0.01 |
| FBW (g) | 6.25±0.14<sup>a</sup> | 5.69±0.62<sup>ab</sup> | 5.15±0.63<sup>ab</sup> | 4.5±0.64<sup>b</sup> | 4.48±0.23<sup>b</sup> |
| BWG (%) | 77.12±3.56<sup>a</sup> | 59.96±16.93<sup>ab</sup> | 45.51±18.63<sup>ab</sup> | 40.79±19.84<sup>ab</sup> | 26.64±6.08<sup>b</sup> |
| SGR (%/day) | 1.9±0.07<sup>a</sup> | 1.53±0.34<sup>ab</sup> | 1.19±0.45<sup>ab</sup> | 1.11±0.47<sup>ab</sup> | 0.78±0.16<sup>b</sup> |
| SR (%) | 100.00±0 | 94.67±5.33 | 100.00±0 | 97.33±2.67 | 100.00±0 |

Data are expressed as mean ± S.E. (N = 3). Different letters refer to $P < 0.05$ . IBW:
initial body weight; FBW: final body weight; BWG: body weight gain; SGR: special
growth rate; SR: survival rate.

### 3.2 Chromium concentration in tissues

Cr accumulation was found in body wall (Fig. 1A), intestine (Fig. 1B) and respiratory tree (Fig. 1C) of sea cucumbers during the experiment. The order of Cr accumulation in tissues was respiratory tree > intestine > body wall after 30 days of dietary Cr^6+^ exposure. The Cr burden in respiratory tree was ∼6 times greater than that in the body wall on day 30.

**Fig. 1.**
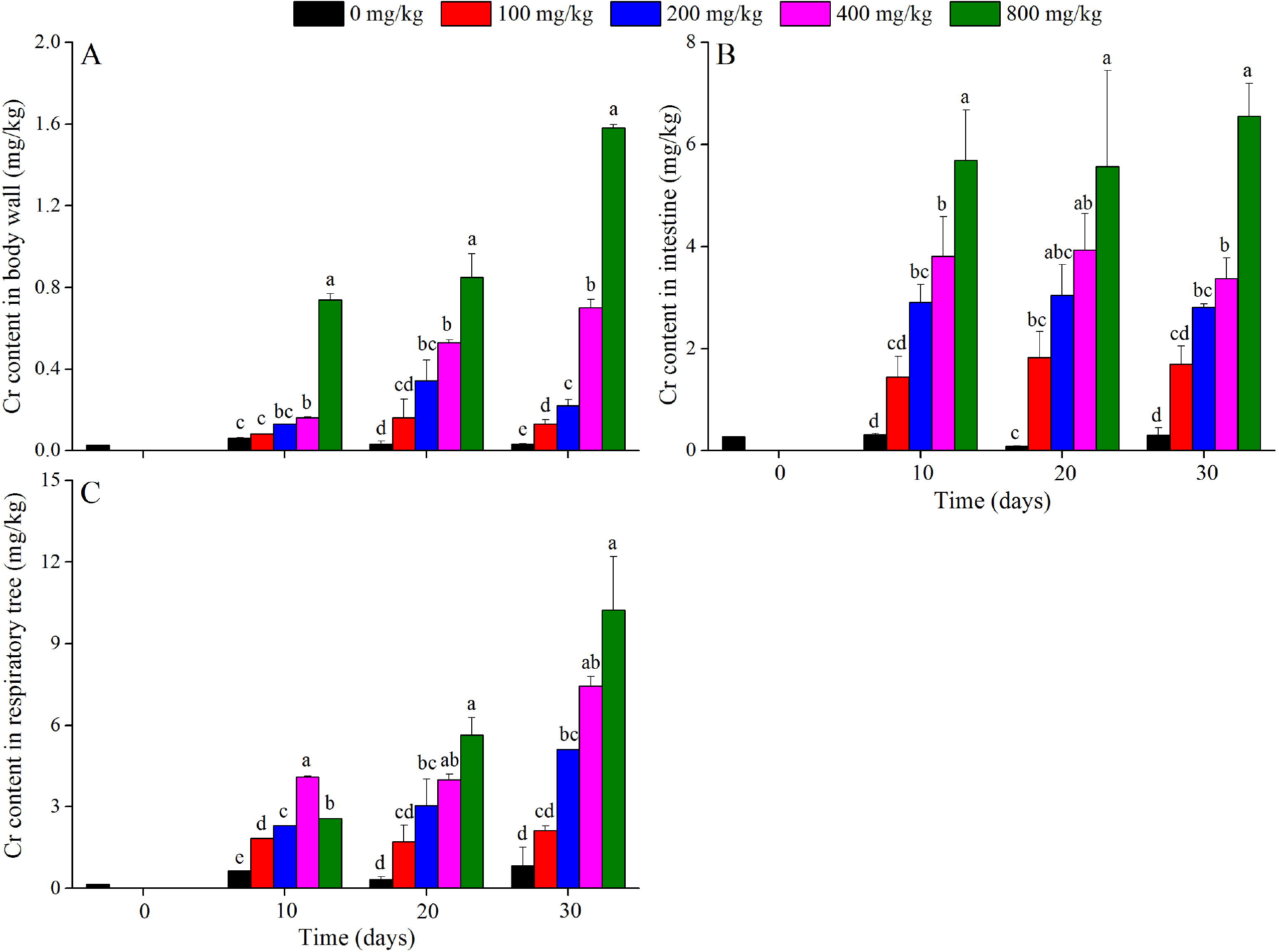
The Cr content in the body wall (A), intestine (B) and respiratory tree (C) of sea cucumbers *Apostichopus japonicus* when they were exposed to Cr^6+^ at different periodsof the experiment (mean ± S.E.). Data at the same exposure times without common letters are considered as *P*□ < 0.05.

The Cr accumulation in body wall increased with concentration increasing. The 800 mg/kg group showed significantly higher Cr burden in body wall on days 10 (0.74 ± 0.03 mg/kg), 20 (0.85 ± 0.11 mg/kg) and 30 (1.58 ± 0.02 mg/kg), compared to that in other groups (*P* < 0.05, Fig. 1A).

The Cr burden in intestine was significantly greater in groups 400 mg/kg (3.81 ± 0.77 mg/kg for day 10, 3.93 ± 0.72 mg/kg for day 20 and 3.37 ± 0.41 mg/kg for day 30) and 800 mg/kg (5.69 ± 0.98 mg/kg for day 10, 5.57 ± 1.87 mg/kg for day 20 and 6.55 ± 0.64 mg/kg for day 30) than that in control group on days 10, 20 and 30 (*P* < 0.05, Fig. 1B).

Significantly higher Cr burden of respiratory tree was found in groups 200 mg/kg (2.30 ± 0.01 mg/kg for day 10, 3.04 ± 0.98 mg/kg for day 20 and 5.09 ± 0.01 mg/kg for day 30), 400 mg/kg (4.09 ± 0.03 mg/kg for day 10, 3.99 ± 0.21 mg/kg for day 20 and 7.45 ± 0.34 mg/kg for day 30) and 800 mg/kg (2.56 ± 0.01 mg/kg for day 10, 5.65 ± 0.63 mg/kg for day 20 and 10.24 ± 1.95 mg/kg for day 30) than that in control group during the experiment (*P* < 0.05, Fig. 1C).

### 3.4 Antioxidant enzyme activities

Significantly lower SOD activity occurred in 400 mg/kg group (129.47 ± 1.35 U/mL) than that in control group (156.94 ± 8.21 U/mL) (*P* □< 0.05, Fig. 2A). The CAT activities in control group (13.91 ± 0.45 U/mL) were significantly higher than those in groups 100 mg/kg (7.05 ± 2.04 U/mL) and 200 mg/kg (10.06 ± 0.74 U/mL) (both *P* □< 0.05, Fig. 2B). Significantly better MDA levels were found in 800 mg/kg group (0.67 ± 0.14 n mol/mL) than those in control group (0.36 ± 0.05 n mol/mL) (*P* □< 0.05, Fig. 2C)

**Fig. 2.**
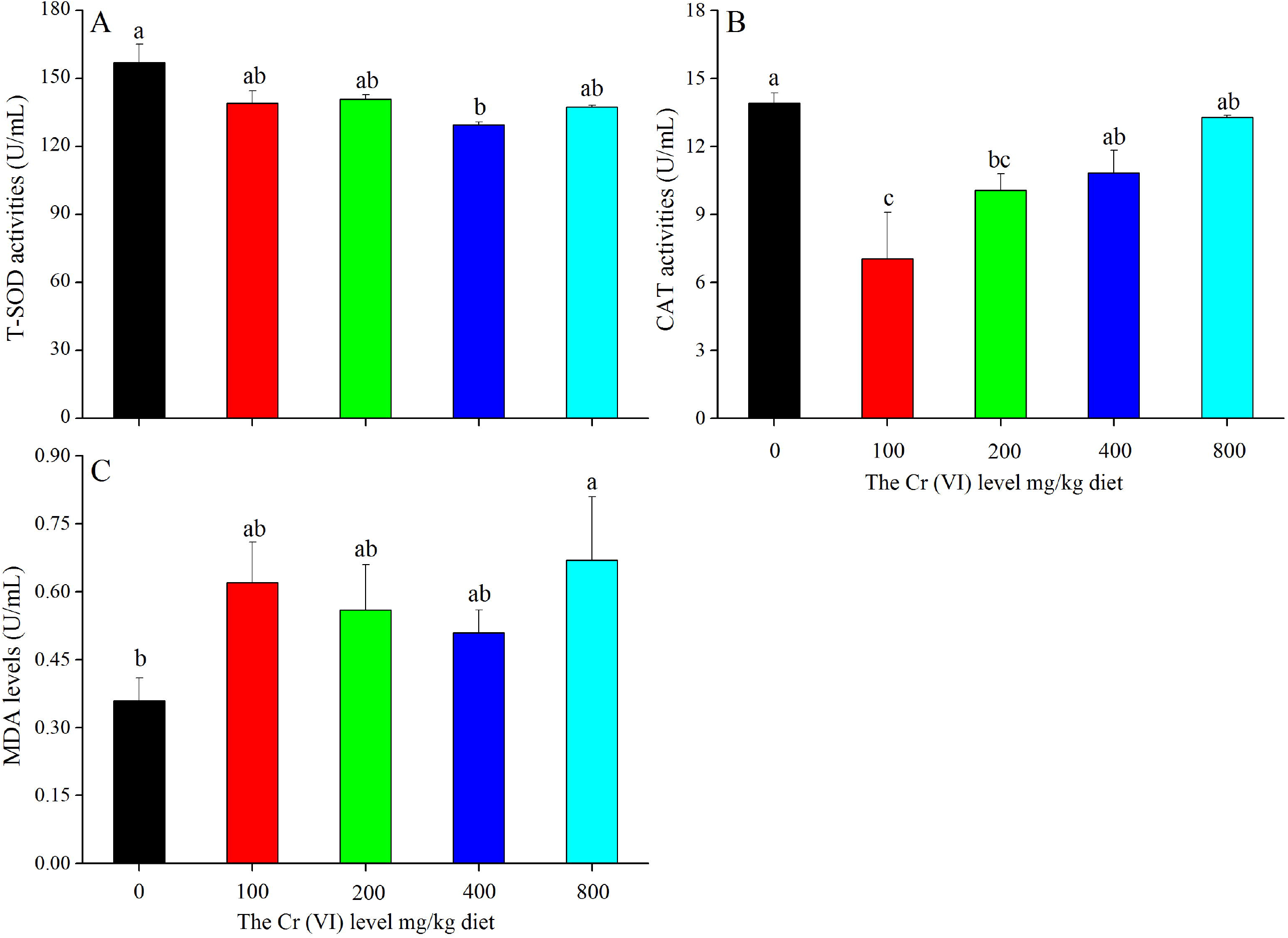
The enzyme activities of SOD (A), CAT (B), MDA (C) of *Apostichopus japonicus* at the end of experiment (mean ± S.E.). Different letters refer to *P* < 0.05.

### 3.4 Hydrolytic enzymes activities

The ACP activities in control group (3.15 ± 0.02 U/100 mL) were significantly higher than those in groups 200 mg/kg (2.72 ± 0.06 U/100 mL), 400 mg/kg (2.28 ± 0.07 U/100 mL) and 800 mg/kg (2.19 ± 0.02 U/100 mL) (*P* < 0.05, Fig. 3A). There were significantly higher ACP activities in 200 mg/kg group than those in groups 400 mg/kg and 800 mg/kg (*P* < 0.05, Fig. 3A). The AKP activities were not significant difference in all groups (*P* □< 0.05, Fig. 3B).

**Fig. 3.**
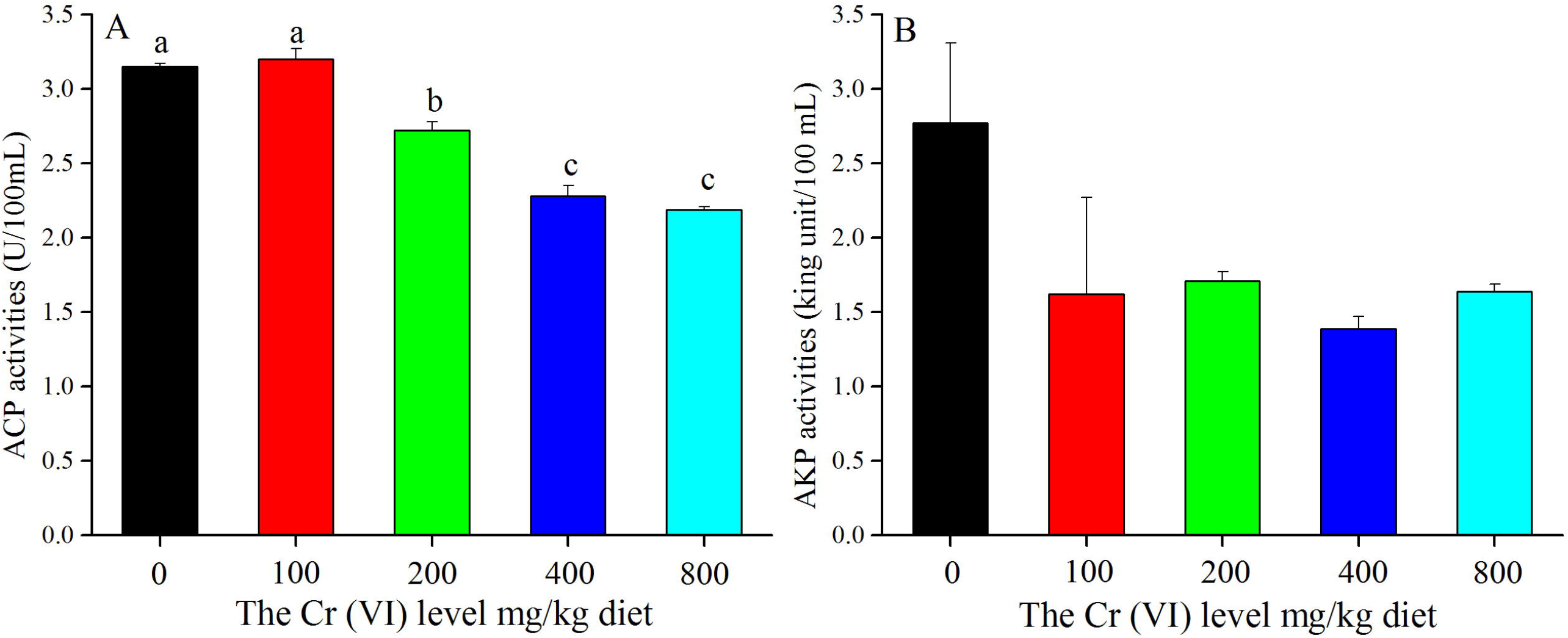
The enzyme activities of ACP (A) and AKP (B) of *Apostichopus japonicus* at the end of experiment (mean ± S.E.). Different letters refer to *P* < 0.05.

## Discussion

Heavy metals are persistent contamination with highly toxic, causing various negative impacts on animals in aquatic ecosystems (Yin et al., 2020). The emission of Cr^6+^ in wastewater, for example, is 24, 844 kg in China in 2017 (Ning, 2020). It is necessary to investigate the effects of chronic dietary Cr^6+^ on bioaccumulation, immunity responses in sea cucumbers *A. japonicus*, especially considering its high commercial and ecological value (Ru et al., 2019).

### 4.1 Chromium concentration in tissues

The bioaccumulation patterns of animals contribute to evaluating environmental metal pollution (Kim et al., 2004). Metal accumulation in aquatic animals (e.g fishes) either directly uptake from the water using their gills or indirectly absorb from diets by intestine (Oost et al., 2003). In this context, dietary exposure is considered as the main cause of metal accumulation in fishes (Hall et al., 1997; Farag et al., 1999) and the intestine is therefore the primary target of metal accumulation (Bernhoft, 2012). For example, Li et al. (2018) observed that the profile of mercury accumulation in tissues was intestine > respiratory tree > body wall. However, the present study showed that the order of Cr accumulation in the experimental tissues was respiratory tree > intestine > body wall, suggesting respiratory tree accumulated the highest concentration of Cr. Respiratory tree of sea cucumbers is similar to gill of fish (Wang et al., 2015), which plays important role in detoxification, gaseous exchange and lon regulation (Goss et al., 1998). Accumulation of Cr in the gills probably is caused by the switch of dietary Cr to the basolateral cell membranes of the gill cells through the bloodstream (Szebedinszky et al., 2001). Wang et al. (2015) documented that the order of tissues Pb accumulation was as follows: body wall > intestine > respiratory tree when *A. japonicus* fed dietary Pb with different levels (100, 500 and 1000 mg Pb/kg dry weight) for 30 days. Li et al. (2019) found that the profile of Cr accumulation in tissues was body wall > intestine after *A. japonicus* was exposed to dietary chromium (58.27 mg Hg/kg dry weight) for 10 days. Together with these previous results, the present study further suggests that the Cr accumulation in tissues is probably related to the exposure time and ability of metal accumulation in each tissue.

### 4.2 Immunity responses

Cr^6+^ enters a cell with the help of sulphur anion transport system (Holland and Avery, 2011) and becomes more stable form (e.g. Cr^3+^) by cellular reductants (Krumschnabel and Nawaz, 2004). Various chemicals, including hydroxyl radicals (OH^-^), superoxide (O^2-^) and hydrogen peroxide (H_2_O_2_), increasing the levels of lipid peroxidation in this process (Vlahogianni et al., 2007). In other words, Cr^6+^ exposure probably directly facilitates the production of reactive oxygen species (ROS) via the Fenton-like cycling mechanism (Sinha et al., 2006) and/or indirectly encourages oxidative stress (OS) by interacting with mitochondria (Pourahmad et al., 2003). ROS attacks cellular constituents (Li et al., 2010; Sturve et al., 2008), alters the defense processes and the antioxidant enzymes activities in cells (Jindal and Handa, 2019). The present study found that significantly higher MDA levels occurred in 800 mg/kg group than those in control group. High MDA levels destroy physiologic integrity by decreasing the resistance and fluidity in cytomembrane (Chandran et al., 2005). These results are consistent with Lushchak et al. (2009), who found that MDA levels increase with the increase of exposed Cr^6+^ concentration (Kubrak et al., 2010) in goldfish, indicating that sea cucumbers were suffering from oxidative stress in relation to ROS generation (Ahmad et al., 2006).

The antioxidant defense system (e.g. SOD and CAT) can neutralize the ROS and consequently protect cells from adverse variation (Ritola et al., 2002; Kubrak et al., 2010). The SOD is considered as the first line against oxygen toxicity in the cells (Li et al., 2010) because it catalyzes the decomposition of superoxide anions into H_2_O and H_2_O_2_ (Shao et al., 2012). The present study found that significantly lower SOD activities occurred in 400 mg/kg group than those in the control group. Significantly decline in SOD activity has also been observed after fishes *Labeo rohita* were exposed to dietary Cr^6+^ (Kumari et al., 2014). Long-term exposure of Cr^6+^ increases the substrate levels (superoxide radicals) of SOD, whereas SOD activity decreased with the increase in its substrate levels in this study, indicating that antioxidant defense probably has been impaired in sea cucumbers. In terms of CAT, as an important enzymatic scavenger (Chen et al., 2015), removing the H_2_O_2_ by catalyzing its decomposition. The present study found that significantly lower CAT activities were found in groups 100 and 200 mg/kg than those in control group. Similar findings also showed that CAT activity decreased significantly in kidney and liver tissues of fishes *Cyprinus carpio* (Kumar et al., 2013) and goldfish (Lushchak et al., 2009). Earlier studies suggested that the overproduction of superoxide anion caused by Cr^6+^ inhibited the CAT activities (Pandey et al., 2001). The glucose-6-phosphate dehydrogenase activity, which is essential for the conservation of CAT activity, probably also affects the CAT activities (Scott et al., 1991). Further, we found that CAT activities in groups 400 and 800 mg/kg increased significantly compared to those in the 100 mg/kg group, while they were not significantly different from those in control group. The interaction between CAT and Cr^6+^ changes the structure of substrate (H_2_O_2_), leading the substrate to more easily reach the enzyme (Zhang et al., 2016). These results suggest higher dietary Cr^6+^ exposure (at least 400 and 800 mg/kg) did not negatively alter the CAT activity, although significantly inhibited CAT activity occurred in lower dietary Cr^6+^ exposure (e.g 100 mg/kg).

It has been well documented that dietary Cr^6+^ exposure influenced immune responses in various animals (Glaser et al., 1985; Arunkumar et al., 2000), including the macrophage functions (Tian and Lawrence 1996), distribution of leukocytes (Arunkumar et al., 2000) and the proliferation of lymphocytes (Wang et al., 1996). The present study further found that ACP activities decreased significantly in groups 200, 400 and 800 mg/kg than those in control group, while no significant differences in AKP activities were found among groups. Jiang et al. (2010) consistently found that significantly decline of ACP occurred in the mud crabs *Scylla paramamosain* after dietary Cr^6+^ exposure. ACP is one of the phosphatases attaching the phosphate groups from other molecules (Cajaraville et al., 2000). It functions when the lysosome fuses with the nuclear endosome (Rajalakshmi and Mohandas, 2005). AKP plays an important role in catalyzing the hydrolysis of phosphate monoesters (Zang et al., 2012). They are both involved in the degradation of lipids and carbohydrates (Ottaviani, 1984; Xue and Renault, 2000). This study found Cr^6+^ displayed an inhibitory effect on ACP activities but not on AKP activities, which probably explains why the nonspecific immune responses were suppressed in fishes *Salmo trutta L*. and *Cyprinus carpio* L. when they were experienced Cr^6+^ exposure (O’Neill, 1981). Collectively, the present study found that high dietary Cr^6+^ exposure (200, 400 and 800 mg Cr^6+^/kg dry weight) probably impairs the immune function of sea cucumbers *A. japonicus*.

## Ethical Approval

All applicable international, national, and/or institutional guidelines for the care and use of animals were followed by the authors.

## Acknowledgments

This work was supported by the National Spark Program of China (2015GA650006). We thank Xiaonan Guan and Xianyu Meng for their assistance.

